# Structural basis for BIRC6 to balance apoptosis and autophagy

**DOI:** 10.1101/2022.12.10.519866

**Authors:** Shuo-Shuo Liu, Tian-Xia Jiang, Fan Bu, Ji-Lan Zhao, Guang-Fei Wang, Guo-Heng Yang, Jie-Yan Kong, Yun-Fan Qie, Pei Wen, Li-Bin Fan, Ning-Ning Li, Ning Gao, Xiao-Bo Qiu

## Abstract

Caspase-9 is the initiator caspase for the intrinsic apoptotic cell death pathway, and is critical to the activation of effector caspases during apoptosis, but how its levels and activities are maintained remains unclear. The gigantic Inhibitor of Apoptosis Protein (IAP) BIRC6/BRUCE/Apollon not only inhibits apoptosis, but also promotes ubiquitination of the key autophagic protein LC3 and inhibits autophagy. Here we show that BIRC6 forms an anti-parallel U-shaped dimer in a 3.6-Å cryo-EM structure with multiple previously unannotated domains, including a ubiquitin-like domain, and discover that the mitochondria-derived pro-apoptotic factor Smac/DIABLO binds BIRC6 by interacting with one BIR domain, two carbohydrate-binding modules and two helices in the central cavity. Notably, Smac outcompetes the effector caspase 3 and the pro-apoptotic protease HtrA2, but not caspase 9, for binding BIRC6. BIRC6 strongly inhibits cellular activity of caspase 9, but weakly suppresses that of caspase 3. Meanwhile, BIRC6 binds LC3 through an LC3-interacting region, probably following dimer disruption of this BIRC6 region. Deficiency in LC3 ubiquitination promotes autophagy and autophagic degradation of BIRC6, and inhibits apoptosis. Moreover, induction of autophagy promotes autophagic degradation of both procaspase-9 and active caspase-9, but not of effector caspases. These results are important to understand how the balance between apoptosis and autophagy is regulated under pathophysiological conditions.

## INTRODUCTION

Apoptosis is a type of programmed cell death (*1*), whereas autophagy primarily supports survival under nutrient-restricted or other stress conditions by removing cargoes including proteins, organelles, and exotic pathogens (*2*). The balance between apoptosis and autophagy is critical for normal development, proper tissue function, and disease pathogenesis (*3*). Apoptosis is executed by activation of the family of caspases (cysteine aspartic acid-specific proteases), which are first synthesized as weakly active zymogens and then proteolytically cleaved to generate active enzymes during apoptosis (*4*). Caspase 9 is the initiator caspase, which activates effector caspases (e.g., caspases 3 and 7), in the intrinsic or mitochondrial apoptotic pathway, where cytochrome c is released from mitochondria to bind Apaf-1 for self-cleavage of procaspase 9 into the active form (*1*). The activity of caspases can be inhibited by the Inhibitor of Apoptosis Proteins (IAPs), which contain one to three tandem baculoviral IAP repeats (BIRs) (*5, 6*). The actions of IAP proteins are counteracted by certain pro-apoptotic factors, especially the mitochondria-derived pro-apoptotic factor Smac/DIABLO. Cytosolic Smac precursor is proteolytically processed in mitochondria into its mature form, which is released into the cytosol to interact with IAPs with its exposed IAP-binding motif (IBM) in response to apoptotic stimuli (*7–9*).

The only essential IAP is the exceptionally large (~530 kDa) membrane-associated protein BIRC6 (also referred to as BRUCE or Apollon), which has a ubiquitin-conjugating (UBC) domain and a single BIR (*10*). BIRC6 promotes ubiquitination and degradation of the mature Smac and effector caspases by serving as both ubiquitin-conjugating enzyme (E2) and ubiquitin ligase (E3) (*11,12*). We demonstrated previously that BIRC6, unlike other IAPs, also binds to the precursor of Smac and promotes its degradation in an IBM-independent manner (*13*). During macroautophagy (referred to below as autophagy), cargoes are sequestered into a double-membrane autophagosome (*14*), which is then fused with the endosome or lysosome to form an autolysosome for cargo degradation (*15, 16*). LC3-I on the phagophore membrane is conjugated to phosphatidylethanolamine to form LC3-II, which is required for the formation of autophagosomes and selective recruitment of cargoes. We showed previously that BIRC6 together with the proteasome activator PA28γ, which usually promotes the ubiquitin-independent protein degradation (*17*), promotes proteasomal degradation of LC3-I and thus inhibits autophagy (*18*). Interestingly, BIRC6 together with the ubiquitin-activating enzyme (E1) UBA6 promotes ubiquitination of LC3 at K51 residue (*19, 20*), though the role of this ubiquitination is unclear.

Although BIRC6 is a gigantic 530 kDa protein, only two small domains, i.e., the 8 kDa BIR domain and the 19 kDa UBC domain, were previously annotated (Fig. 1A) (*11*). It is unknown how BIRC6 interacts with such a variety of binding proteins and how BIRC6 utilizes its ubiquitination function to regulate both apoptosis and autophagy. Here we reveal multiple domains in the 3.6-Å cryo-EM structure of BIRC6, which forms an anti-parallel U-shaped dimer, in addition to the two sets of BIR and UBC located next to each other, and provide evidence how the balance between apoptosis and autophagy is regulated.

**Fig. 1.**
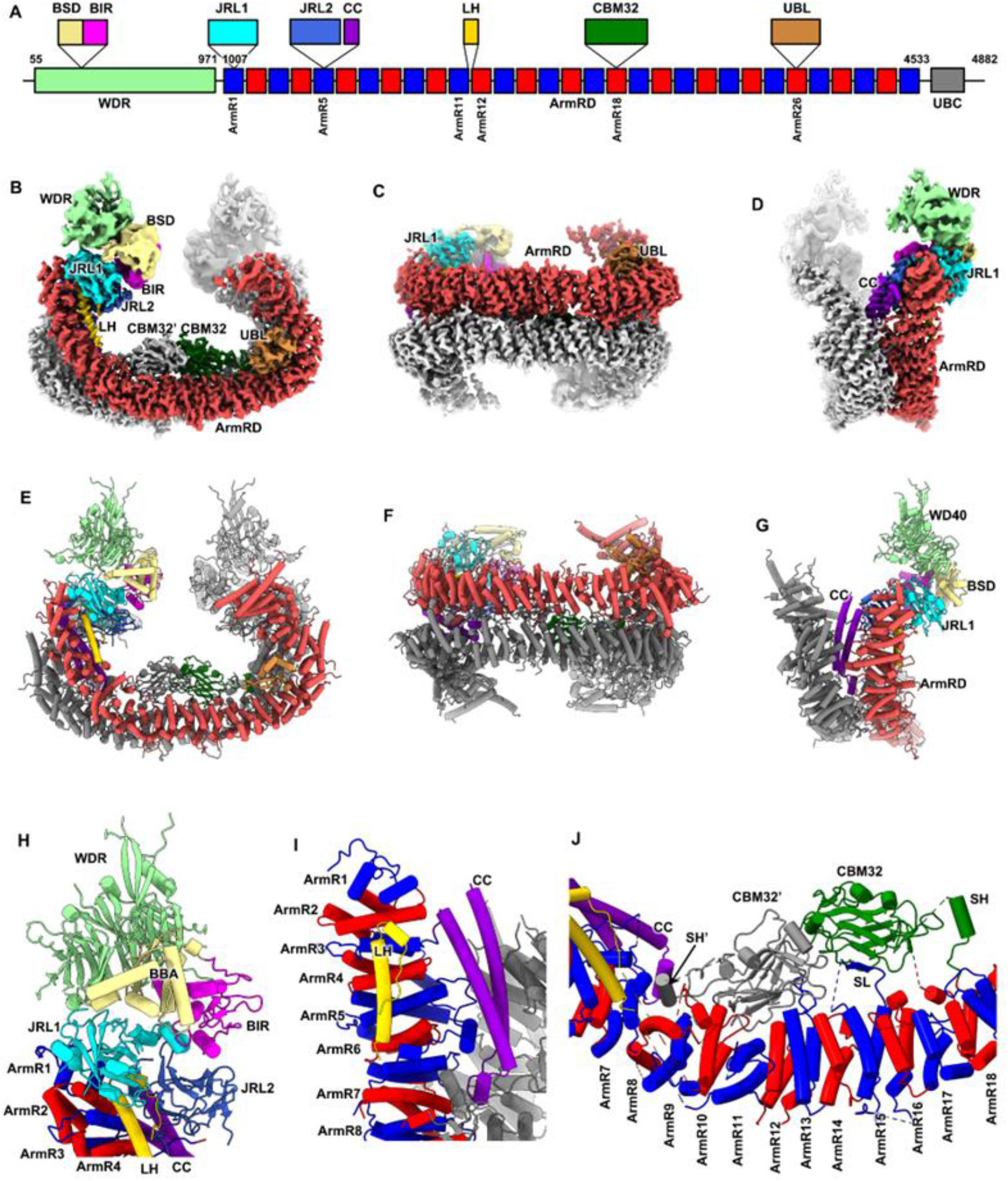
BIRC6 forms an anti-parallel U-shaped dimer with multiple protein-binding domains. (**A**) Schematic domain organization of mouse BIRC6. The 31 ArmRs are colored blue and red alternatively. (**B** to **D**) Cryo-EM structure of BIRC6 dimer displayed in three different views. For one monomer, ArmRD are colored valentine red, and the other domains are color-coded as that in (**A**) The other monomer is colored gray. (**E** to **G**) Same as (**B** to **D**) but shown in cartoon representation. (**H**) Zoomed-in view of the boxed region in (**E**) but with a slight rotation. (**I**) Zoomed-in view of the boxed region in (**G**) but with a 180°-rotation. (**J**) Zoomed-in view of the boxed region in (**E**) but with a slight rotation. The domains in (**H** to **J**) are colored-coded as that in (**A**).

## RESULTS

### BIRC6 forms dimer with multiple domains

To characterize the structure of BIRC6 and to understand the structure-function relationship, we expressed and purified the full-length mouse BIRC6 using HEK293F cells, and determined the cryo-EM structure of BIRC6 at a global resolution of 3.6 Å (fig. S1 and S2, and Table S1). The core regions were well resolved at side-chain resolution, allowing an accurate sequence assignment during modeling (Fig. 1, B to D, and fig. S2, E and F).

The overall structure of BIRC6 is organized as a symmetric homodimer in a U-shape (Fig. 1, B to G), consistent with the elution profile from gel filtration (fig. S1). The two monomers interact with each other side-by-side in an antiparallel fashion (Fig. 1, B to G), forming a very extensive dimer interface. The central section of BIRC6 is responsible for dimer interactions mediated by an armadillo-repeat domain (ArmRD), which is composed of 31 armadillo repeats (ArmR) (Fig. 1, A to G, and fig. S3A). Unlike typical armadillo repeats, ArmRD of BIRC6 contains many insertion sequences located either in an inter-ArmR or intra-ArmR manner. Most of these insertions are flexible and not fully resolved in the map. Several insertion domains or motifs, due to their participation in the inter- or intra-molecular interaction, are stabilized and could be modeled unambiguously (Fig. 1, A to G, and fig. S3, C to L).

Within ArmR1, an insertion forms a β-sandwich motif with its core folded as a “beta-jelly roll” topology (Fig. 1, A and H, and fig. S3, C and D). Since no significant structural homology was identified using the DALI server (*21*), this insertion was termed jelly-roll like domain 1 (JRL1) hereafter. Similarly, another jelly-roll like domain (JRL2) is inserted in ArmR5 (Fig. 1, A and H, and fig. S3, E and F), intermediately followed by a coiled-coil motif (CC) (Fig. 1, A, D, G and I, and fig. S3G). Next, a long helical motif (LH) inserted between ArmR11 and ArmR12 interacts tightly with ArmR2-7, JRL1 and JRL2 of the same monomer, contributing to the stabilization of JRL1 and JRL2 (Fig. 1, A, H and I, fig. S3H). Further towards the C-terminus, a third jelly-roll like domain is inserted in ArmR18 (Fig. 1, A and I). DALI search indicates that it is a domain of carbohydrate-binding module family 32 (CBM32) (fig. S3, I and J). The CBM32 has no contact with the ArmRs from the same BIRC6 monomer, except that a short loop (SL) from ArmR15 stretches to form an additional β-strand to complement one β-sheet of CBM32 (Fig. 1J).

At the C-terminal region of ArmRD, a ubiquitin-like domain (UBL) inserted in ArmR26 makes extensive interactions with ArmR20-21 and ArmR24-25 (Fig. 1, A, B, and E, and fig. S3A, K and L). Consequently, these interactions with the UBL generate a sharp bending around ArmR21-24, leading to the formation of the U-shape of the global structure (fig. S3A). In addition, the E1 UBA6, which can bind ubiquitin, was highly enriched in the BIRC6 complex (fig. S1E), raising the possibility that UBA6 directly binds BIRC6 through this ubiquitin-like domain.

The major component of the N-terminal region is a WD40 repeat (WDR) β-propeller domain (Fig. 1, A and H, and fig. S3B). There is a long insertion between the second and the third blades of the WDR, which folds into two structural domains, BIR and a tightly associated domain (Fig. 1A, and fig. S3B). This domain mediates the indirect interaction between BIR and WDR and thus termed BIR-stabilizing domain (BSD) hereafter (Fig. 1A, and fig. S3B). All the three domains, WDR, BSD and BIR, interact with the JRL1 domain, orienting the BIR domain toward the center of the U-shape structure (Fig. 1H). The N-terminal section of BIRC6 is dynamic relative to the central section (fig. S2G), which might be beneficial for its substrate capturing. The C-terminal UBC domain (Fig. 1A) is highly flexible and completely invisible in the density map. Considering that the C-terminal end of ArmRD is positioned close to the central space of the U-shaped structure, the UBC domain could be close to the BIR domain (Fig. 1, B and E, and fig. S3A).

The dimerization of BIRC6 is mediated by the two ArmRDs exclusively. There are extensive side-by-side interactions between aligned ArmRs from the two monomers. In addition, two ArmR insertions also participate in the dimerization (Fig. 1, C and F). One is CBM32 inserted in ArmR18, which forms extensive interactions with the ArmR7-13 and CBM32 of the other monomer (Fig. 1J). The other is the CC inserted in ArmR5, which is sandwiched between the region of ArmR1-6 of one monomer and ArmR22-27 of the other to bridge the interaction between the N-terminal and C-terminal segments of two BIRC6 monomers (Fig. 1, D, G and I).

In summary, the cryo-EM structure reveals the domain organization of BIRC6 and a central space for potential substrate accommodation. This anti-parallel arrangement of two monomers would bring close the two sets of functional domains (such as BIR and UBC), suggesting that BIRC6 dimer could work either *in cis* or in trans to fulfill its role in substrate ubiquitination.

### Mechanisms by which Smac binds BIRC6

During image processing, we noticed some residual unassigned densities above the two CBM32 domains (fig. S2, C and D). Considering that both BIR and UBC domains are facing the central cavity, the residual density might be from endogenous substrates of BIRC6. With several rounds of local 3D classification on this region, several classes with strong additional density above two CBM32 domains were obtained. Refinement of one class led to a 4.7-Å density map in which the extra density was resolved partially at secondary structural level. Mass-spectrometric analysis of our BIRC6 samples indicated the presence of Smac as a co-purified substrate (fig. S1E). Accordingly, a rigid-body fitting of the dimeric Smac complex (Fig. 2D) (*22*) reports a good match between the model and the density. In addition to the density of the Smac dimer, two additional single α-helices (termed unknown helix, UNH) with an apparent C2 symmetry could be found to stack on the Smac dimer (Fig. 2, A to E). The two helices might each originate from one BIRC6 monomer, and given their binding position, they might be important to promote the binding of Smac.

**Fig. 2.**
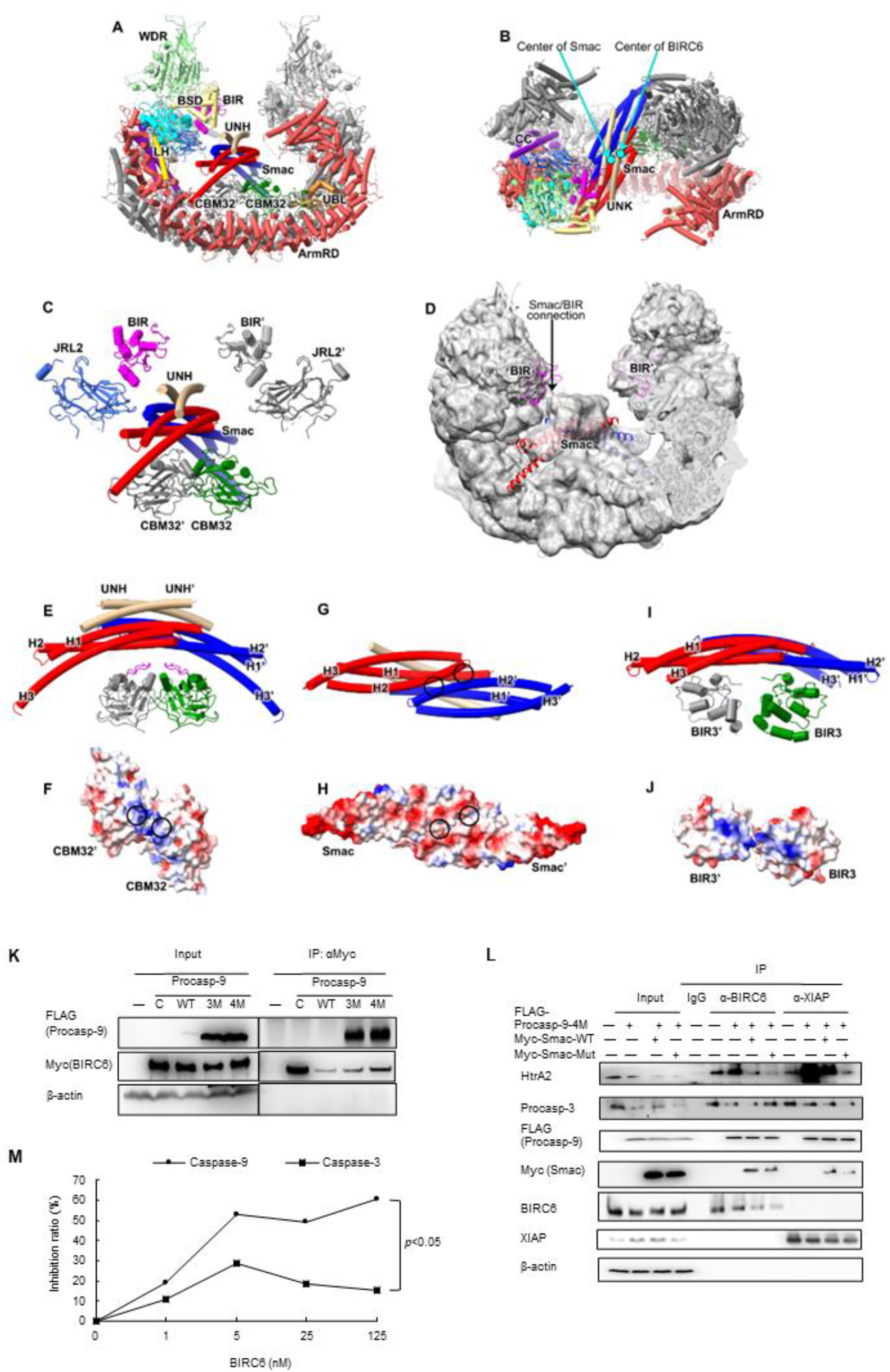
Smac binds BIRC6 dimer, and outcompetes caspase 3 and HtrA2, but not caspase 9. (**A** and **B**) Global structure of Smac-BIRC6 complex displayed in two different views in cartoon representation. The domains of BIRC6 are color-coded as that in Fig. 1, B to G. The two monomers of Smac are colored red and blue, respectively. The two UNHs are colored brown. The two cyan dots in panel Fig. 2B indicate the centers of Smac dimer and BIRC6 dimer, respectively. (**C**) Structure of Smac-BORC6 complex showing only Smac and adjacent domains BIRs, JRL2s and CBM32s of BIRC6 dimers. The domains or subunits are color-coded as that in Fig. 2. (**D**) Cryo-EM density map of Smac-BIRC6 complex displayed in low counter level, and with atomic model superimposed. The two Smac monomers are colored red and blue, respectively. The two BIRs are colored magenta. The density connecting Smac and BIR is indicated. (**E**) Structure of Smac interacting with CBM32, shown in cartoon representation. The domains or subunits are color-coded as that in Fig. 2, A and B. The loops in the two CBM32s interacting with Smac are colored magenta. UNHs and the helices of Smacs are labeled as indicated. (**F**) The electrostatic potential surface of the CBM32 dimer. The two sites contacting Smac are indicated by black circles. (**G**) Structure of Smac dimer shown in cartoon representation, in a view with a 90°-rotation to that in panel (**E**). The sites contacting CBM32s are indicated by black circles. (**H**) Same as **G**, but showing the electrostatic potential surface of the Smac dimer. (**I**) Structure of the dimeric Smac-XIAP-BIR3 complex by aligning the Smac in monomeric Smac-XIAP-BIR3 complex to Smac in this study. (**J**) The electrostatic potential surface of the two XIAP-BIR3s, displayed in a view with a 90°-rotation to that in panel (**I**) to show the electrostatic potential of the interface to Smac. (**K**) HEK293T cells were co-transfected with Myc-BIRC6 and the FLAG-tagged wild-type (WT), triple mutant (E306A–D315A–D330A; Procasp 9-3M) or quadruple mutant procaspase 9 (E306A–D315A–D330A-C287A; Procasp 9-4M). Immunoblotting was performed following immunoprecipitation with anti-Myc antibodies. (**L**) HEK293T cells were transfected with the FLAG-tagged quadruple mutant procaspase 9 (Procasp 9-4M) in combination with Myc-tagged wild-type or mutant Smac as indicated. Immunoblotting was performed following immunoprecipitation with anti-BIRC6 or XIAP antibodies in the presence of 50 μM of the pan-protease inhibitor Z-VAD-fmk. (**M**) Active caspases in HEK293T cell extracts were incubated with the purified FLAG-tagged BIRC6 at indicated concentrations. The caspase activities were analyzed using the caspase 9 substrate Ac-LEHD-AMC or caspase 3 substrate Z-DEVD-7-AMC. The release of AMC from the substrates was monitored continuously at 380/460 nm (excitation/emission) at 30 °C for 30 min. Data in Fig. 2, K to M are representative of one experiment with three independent biological replicates.

We noticed that the Smac dimer is slightly off-axial with the BIRC6 dimer (Fig. 2, A to C). Smac dimer leans to one side of the central cavity and close to the BIR and JRL2 domains of this side (Fig. 2, A to C). Smac usually binds to the BIR domain of other IAPs with the N-terminal IBM to antagonize the inhibition of IAP to caspases (*23*). Although the interaction of Smac-IBM with BIR could not be identified due to the resolution limitation, the density connection between Smac and one BIR domain could be observed when the density map was shown in lower contour level (Fig. 2D). In contrast, the Smac connection to the other BIR domain was much weaker (Fig. 2D), indicating a potentially complex regulation of Smac on the two BIR domains.

The stabilization of Smac is mainly dependent on the interactions with the CBM32 dimer, which is mediated by the helices H2/H2’ of Smac dimer and the short loops (residues 3209-3219) of the CBM32 dimer in electrostatic interactions (Fig. 2, A, C, and E to H). The regions of Smac contacting the two CBM32 domains are slightly different, due to the symmetry mismatch between BIRC6 and Smac (Fig. 2, G and H). In addition to the principal interaction of Smac-IBM with the BIR domain, another interface between Smac and BIR3 of XIAP was previously also reported (*23*). By structural superimposition, it is surprising to find that this second interface is similar to that of Smac-CBM32 interaction and both are mediated by electrostatic interactions (Fig. 2, I and J). CBM32 might act as a functional equivalent of BIR3 of XIAP to build the second Smac interface, which could explain why BIRC6 has only one BIR domain while XIAP has three (*24*).

This binding pattern of Smac shows that the central space of the BIRC6 dimer is the substrate binding pocket. Indeed, some other unassigned residual densities were also observed in the central space of the non Smac-binding groups (fig. S2H). Given that various domains and insertions are facing this cavity, BIRC6 ought to have a complex regulation on the recognition of its substrates.

### Smac cannot reduce caspase 9 binding

We showed previously that BIRC6 directly inhibits the activity of the purified caspase-9, but not the purified effector caspase caspase-3 (*13*). Although BIRC6 binds the purified processed caspase 9 (*13*), it remains unclear whether it associates with procaspase 9 in cells. Since the transfected wild-type procaspase 9 was quickly processed (Fig. 2K), we constructed the non-cleavable procaspase-9 triple mutant (E306A–D315A–D330A; Procasp9-3M) and the quadruple mutant with additional active site mutation (E306A–D315A–D330A–C287A; Procasp9-4M). Transfected Myc-BIRC6 could co-immunoprecipitate with both triple and quadruple mutants of FLAG-procaspase 9 (Fig. 2K). Accordingly, the purified BIRC6 could partially co-elute with His-procasp9-4M in the gel filtration assay (fig. S4A). Overexpression of the Myc-tagged wild-type Smac did not affect the association of BIRC6 or XIAP with the FLAG-tagged quadruple mutants of procaspase 9, but markedly reduced their association with the pro-apoptotic protease HtrA2/Omi or procaspase 3 (Fig. 2L), hinting that BIRC6 or XIAP binds procaspase 9 in a mechanism different from that for HtrA2 or procaspase 3. Moreover, the Smac with IBM mutations, where the AVPI stretch was mutated into AAAI, further reduced the co-immunoprecipitation of either BIRC6 or XIAP with HtrA2 (Fig. 2L), suggesting that the region other than the IBM of HtrA2 can also mediates its association with BIRC6. In contrast, this Smac mutant did not affect the co-immunoprecipitation of BIRC6 with procaspase 3, though further reducing the co-immunoprecipitation of XIAP with procaspase 3 (Fig. 2L), suggesting that BIRC6 associates with procaspase 3 in a manner different from XIAP. We then showed that the purified BIRC6 could strongly inhibit the activity of caspase 9, but only weakly for caspase 3, in the HEK293T cell lysates following caspase activation by cytochrome c and dATP (Fig. 2M, and fig. S4, B to D). Thus, Smac outcompetes caspase 3 and HtrA2, but not caspase 9, for binding BIRC6 in cells, resulting in strong inhibition of the initiator caspase 9 and weak inhibition of the effector caspase 3 by BIRC6.

### Caspase 9 can be degraded by autophagy

Nutrient starvation and rapamycin inhibit the activity of mTOR, and induce autophagy (*25*). Starvation in Hanks’ Balanced Salt Solution (HBSS) or treatment with rapamycin reduced the levels of procaspase-9, but not of procaspase-3 and procaspase-7, in HEK293T (Fig. 3A) or MDA-MB-453 cells (fig. S5A). The levels of procaspase-9 decreased gradually upon starvation in COS7 cells, especially at 4 h, and were reversed by the autophagy inhibitor Bafilomycin A1 (Baf A1) (Fig. 3B). Moreover, the levels of active caspase-9, though very low, also decreased upon starvation and could be reversed by Baf A1 (Fig. 3B). In the HEK293T cells transfected with Myc-procasp9-3M, the half-life of the Myc-procasp9-3M was about 12 h, but was reduced to less than 8 h in the presence of rapamycin (fig. S5B). Atg5 is essential for the formation of autophagosome (*26*). The treatment with rapamycin reduced the levels of procaspase-9 in the wild-type mouse embryonic fibroblast (MEF) cells, but not in the Atg5-deficient cells (Fig. 3C). Little caspase-9 co-localized with LC3-II under normal condition, but its co-localization with LC3-II increased dramatically upon starvation (Fig. 3D). These results suggest that both procaspase 9 and active caspase 9 are degraded by autophagy upon autophagic stimulation.

**Fig. 3.**
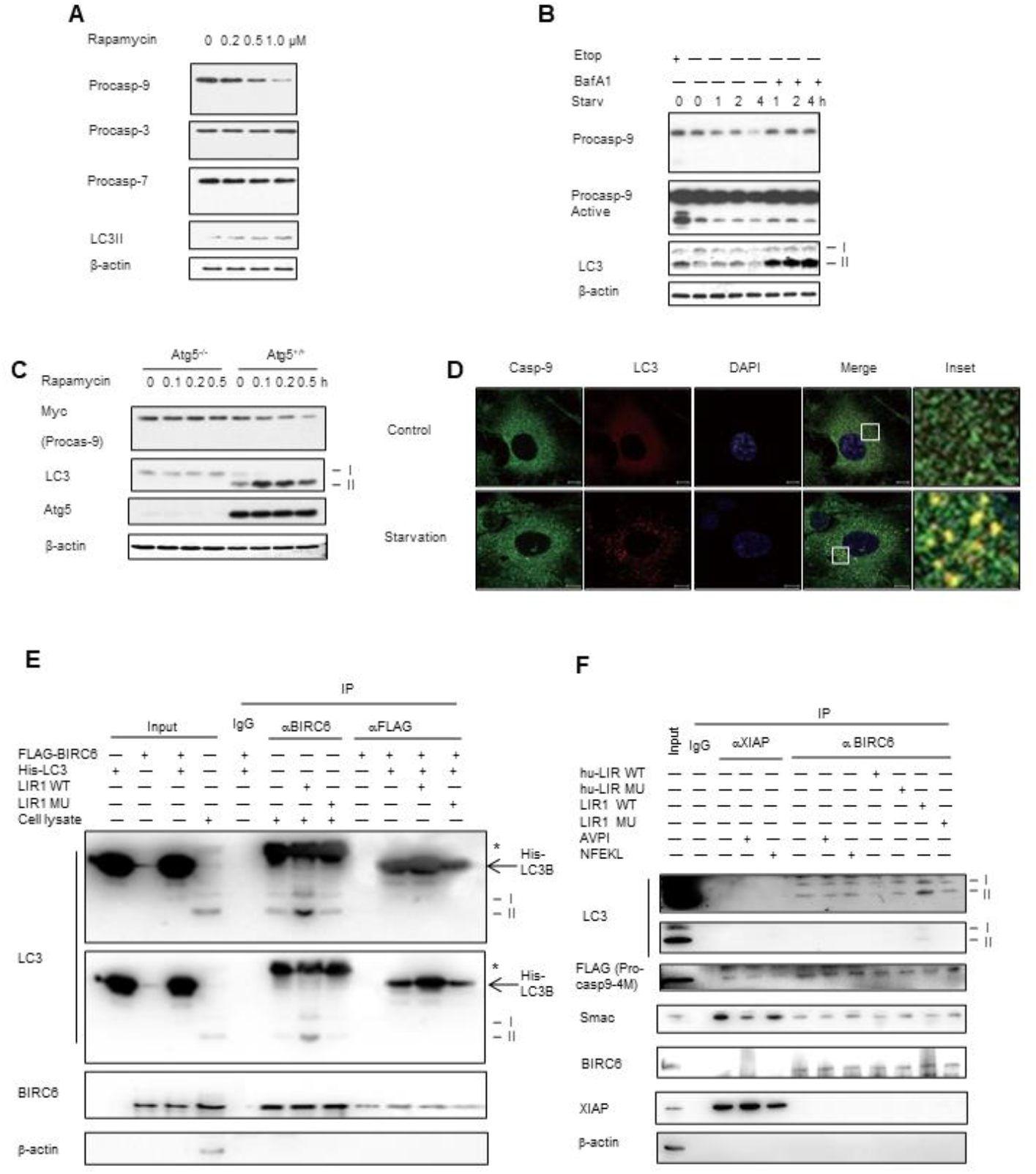
Procaspase 9 can be degraded by the autophagy mediated by LC3, which binds the LC3-interacting region of BIRC6. (**A**) MDA-MB-453 cells were treated with rapamycin at 0, 0.2, 0.5, or 1.0 μM for 24 h. Protein levels were analyzed by immunoblotting. (**B**) COS7 cells were starved in HBSS with or without 10 nM of Baf A1 for 0, 1, 2, or 4 h. Treatment with 20 μM of etoposide for 24 h was served as a control for caspase activation. Protein levels were analyzed by immunoblotting. (**C**) Atg5^+/+^ and Atg5^-/-^ MEF cells were transfected with Myc-procaspase 9 and then treated with rapamycin at the indicated concentration (μM) for 24 h. The protein levels were detected by immunoblotting. (**D**) COS7 cells were starved in HBSS medium for 0 or 2 h, and then subjected to immunostaining and confocal microscopy. DNAs in nuclei were stained with DAPI. n = 30 images, Scale bar: 10 μm. The white rectangles indicate the enlarged areas. (**E**) BIRC6 was immunoprecipitated from the HEK293T cell lysates by the anti-BIRC6 antibody or from the mixture of the purified His-LC3B and FLAG-BIRC6 by anti-FLAG, following supplementation of the LIR1 peptide (NPQTSS<u>F</u>LQ<u>V</u>LV) or its mutant (NPQTSS<u>A</u>LQ<u>A</u>LV). Protein levels were analyzed by immunoblotting. Asterisks represent IgG light chains from anti-BIRC6 antibodies. (**F**) BIRC6 was immunoprecipitated from the lysates of HEK293T transfected with the FLAG-procasp9-4M by the anti-BIRC6 or anti-XIAP antibody, following supplementation of LIR1 (NPQTSS<u>F</u>LQ<u>V</u>LV), hu-LIR (EPLDFDWEIVLEEEM), AVPI, NFEKL, or the related mutant LIR peptides. Protein levels were analyzed by immunoblotting. Data are representative of one experiment with three independent biological replicates.

### BIRC6 binds LC3 through LIR

Because BIRC6 associates with both caspase 9 and LC3, the autophagic degradation of caspase 9 could potentially be mediated by BIRC6. Thus, we attempted to explore the mechanisms by which BIRC6 interacts with LC3. Proteins usually bind LC3 through a LC3-interacting region (LIR) with the consensus sequence, X-3X-2X-1[W/F/Y]X1X2[L/I/V]X4X5, where alternative letters are placed in square brackets (*27*). We showed previously that BIRC6 contains a putative LIR (amino acids *4*670-NPQ TSS <u>F</u>LQ<u>V</u> LV-4681 in mouse) and co-immunoprecipitates with LC3 (*18*). Unexpectedly, addition of this putative LIR peptide (i.e., LIR1, NPQTSS<u>F</u>LQ<u>V</u>LV) could not reduce the precipitation of LC3 with BIRC6 (Fig. 3E). Conversely, the LIR1 peptide, but not its mutant (NPQTSS<u>A</u>LQ<u>A</u>LV, where the replaced residues are underlined), dramatically increased the amounts of both LC3-I and LC3-II associated with BIRC6 (Fig. 3E). To clarify whether this effect is direct, we purified the precursor of LC3-I (i.e., LC3B), which was expressed in bacteria. When incubated with the purified FLAG-tagged BIRC6, the LIR1 peptide still greatly increased the association of LC3B with BRIC6 in vitro (Fig. 3E, and fig. S5, C and D). Next, we synthesized a <u>h</u>ypothetical universal LIR peptide with a sequence different from any known protein (hu-LIR, EPLDFD<u>W</u>EI<u>V</u>LEEEM), which theoretically competes with any LIR motif (*28*). The hu-LIR peptide, but not its mutant (EPLDFD<u>A</u>EI<u>A</u>LEEEM, where the replaced residues are underlined), could reduce the association of LC3 with BIRC6 (Fig. 3F). In contrast, the AVPI stretch of Smac or an unrelated peptide could not affect binding of LC3 to BIRC6, though the AVPI stretch could reduce the association of Smac with XIAP (Fig. 3F). Because BIRC6 forms a dimer as illustrated above, we reason that the dimerization of BIRC6 usually blocks binding of its LIR to LC3, whereas the certain cellular signal (e.g., apoptotic or excessive autophagic signal) or the LIR1 peptide might interfere with the dimerization of this region so that the originally buried binding surface of the BIRC6 LIR becomes exposed for binding to LC3, leading to eventual degradation of LC3 by the proteasomal or autophagic pathway.

### LC3 ubiquitination promotes apoptosis

We showed previously that monoubiquitination of the adaptor protein SIP/CacyBP can switch the cell status between apoptosis and autophagy, likely by controlling degradation of BIRC6 (*18*). To test whether the BIRC6-mediated ubiquitination of LC3 balances apoptosis and autophagy, we constructed a LC3 mutant (K51R) in which the only ubiquitination site K51 was mutated into R. This mutation inhibited degradation of the GFP-fused LC3 in HeLa cells treated with the translation inhibitor cycloheximide (CHX) (Fig. 4A). Overexpression of the proteasome activator PA28γ facilitated degradation of the wild-type LC3, but not its K51R mutant, in either HeLa or HEK293T cells (fig. S6, A and B). However, the K51R mutation did not affect the association of LC3 with PA28γ as shown by the co-immunoprecipitation assays (fig. S6C). The K51R mutation markedly increased the ratio of LC3-II/I, a marker for the occurrence of autophagy, in HEK293T cells treated with Baf A1 in conjunction with rapamycin (Fig. 4B). Accordingly, this mutation promoted autophagic degradation of BIRC6 and the autophagy receptor protein p62 in the cells treated with rapamycin (Fig. 4C). Further, the K51R mutation led to formation of much more autophagosomes, as marked by LC3 puncta, in HeLa cells (Fig. 4D). On the other hand, the LC3 mutation at K51R obviously reduced the levels of the cleaved caspase 3 and apoptosis in response to treatment of the topoisomerase inhibitor etoposide (Fig. 4, E and F). These results suggest that the BIRC6-mediated ubiquitination of LC3 usually inhibits autophagy, and promotes apoptosis.

**Fig. 4.**
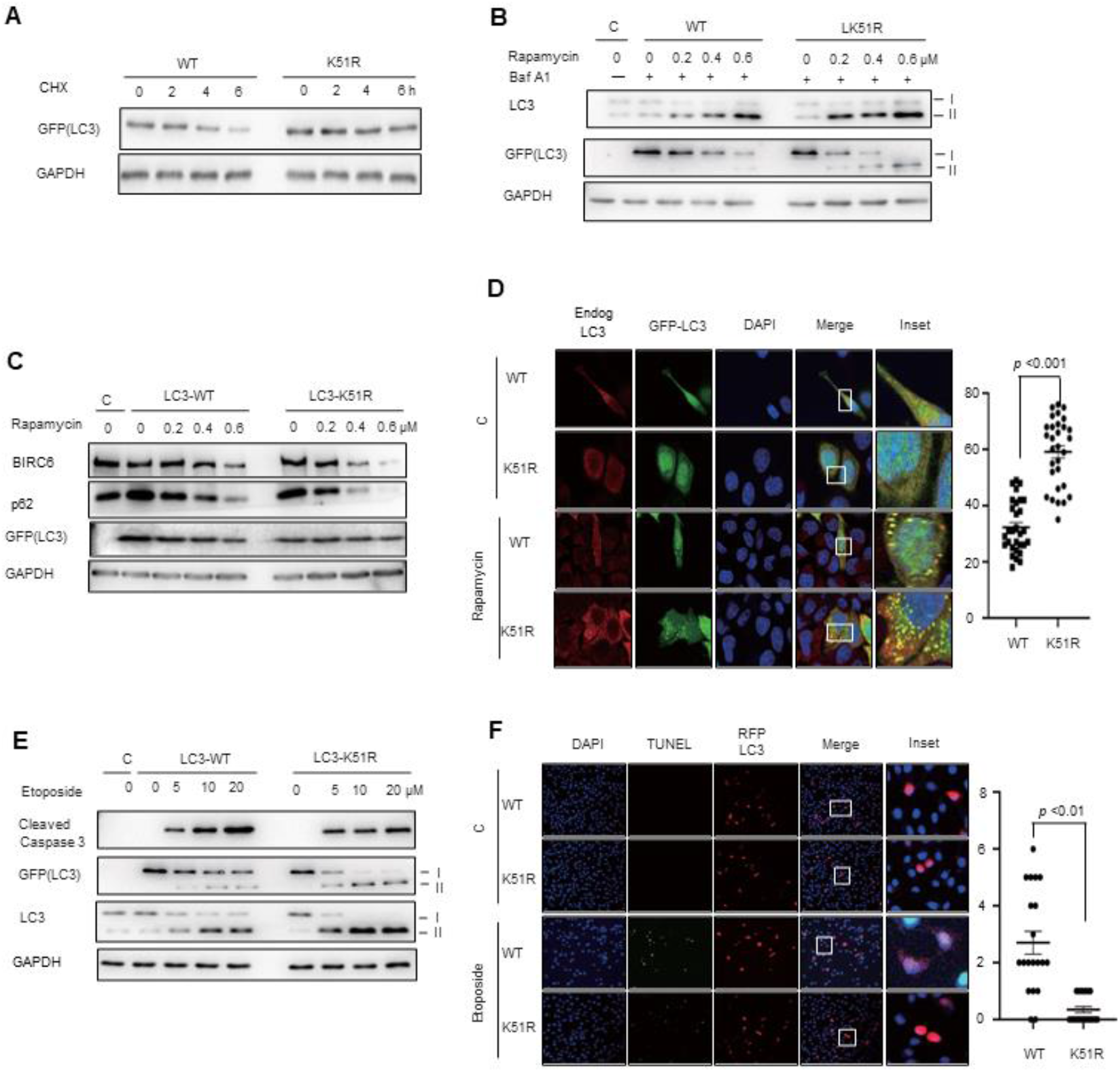
Deficiency in LC3 ubiquitination promotes autophagy, but suppresses apoptosis. (**A**) HEK293T cells were transfected with GFP-LC3-WT or LC3-K51R and then treated with 100 μg/mL CHX for the indicated periods of time. Protein levels were analyzed by immunoblotting. (**B**) HEK293T cells were transfected with GFP-LC3-WT or LC3-K51R and then treated with rapamycin (0, 0.2, 0.4, or 0.6 μM) for 24 h and/or 0.1 μM Bafilomycin A1 for 2 h. Protein levels were analyzed by immunoblotting. (**C**) HEK293T cells were transfected with GFP-LC3-WT or LC3-K51R and then treated with rapamycin (0, 0.2, 0.4, or 0.6 μM) for 24 h. Protein levels were analyzed by immunoblotting. (**D**) HeLa cells were transfected with GFP-LC3-WT or LC3-K51R and then treated with 0.5 μM rapamycin for 24 h. The numbers of LC3 puncta were detected under a confocal microscope following immunostaining. n = 30 images, Scale bar: 10 μm. (**E**) HeLa cells were transfected with GFP-LC3-WT or LC3-K51R and then treated with etoposide (0, 5, 10, or 20 μM) for 24 h. Protein levels were analyzed by immunoblotting. (**F**) HeLa cells were transfected with RFP-LC3-WT or LC3-K51R and then treated with 20 μM etoposide for 12 h. The numbers of apoptotic cells were detected by TUNEL assay under a confocal microscope. n = 30 images, Scale bar: 10 μm. Data are representative of one experiment with three independent biological replicates.

## DISCUSSION

An appropriate level of caspase 9 is the key to determine cell death or survival, but how the levels of caspase 9 are maintained remains unclear. The results in this study strongly suggest that the only essential IAP BIRC6 plays a vital role in inhibiting apoptosis by suppressing caspase 9 under normal conditions (fig. S6D). We determine the 3.6-Å cryo-EM structure of BIRC6, which forms an anti-parallel U-shaped dimer with the two sets of functional domains BIR and UBC located next to each other. Unexpectedly, Smac was highly enriched in the BIRC6 complex in regularly cultured cells (fig. S1E). A dimer of Smac was found located in the central cavity of a subpopulation of the particles directly purified from cells. BIRC6 binds Smac through one BIR domain, two helices and a domain of carbohydrate-binding module family 32 (CBM32) in a mechanism different from any other IAPs.

Since overexpression of Smac could not exclude caspase 9 from binding cellular BIRC6, the primary role of Smac in apoptosis is most likely to release pro-apoptotic protease HtrA2 and the effector caspases, all of which were enriched in the BIRC6 complex (fig. S1E), from BIRC6 binding. Failure of Smac to release procaspase 9 from BIRC6 presumably allows the sufficient chance for BIRC6 to promote ubiquitination and proteasomal degradation of procaspase 9 in addition to inhibiting caspase 9 activity directly. This notion is in accord with the fact that the E1 UBA6 was highly enriched in the BIRC6 complex (fig. S1E). Because an E1 usually binds ubiquitin, whether UBA6 directly binds the ubiquitin-like domain of BIRC6 deserves further investigation.

BIRC6 might target caspase 9 for autophagic degradation in response to autophagic stimuli. Both procaspase 9 and active caspase 9 underwent autophagic degradation upon rapamycin treatment or nutrient starvation. Coincidently, knockdown of the autophagic receptor p62 markedly increases the levels of active caspase 9 and active caspase 3 in the Cadmium-treated JEG-3 trophoblast cells (*29*). Unlike autophagic degradation of active caspase 8 or active caspase 3 in response to apoptotic stimuli (*30, 31*), the degradation of the initiator caspase (i.e., caspase 9) during autophagy might hold the effector caspases, such as caspases 3 and 7, in inactive forms, supporting that autophagic degradation of caspase 9 plays a more important role in inhibiting apoptosis under physiological or pathological conditions.

In response to apoptotic stimuli, BIRC6, together with UBA6, should promote ubiquitination of LC3 at K51, leading to inhibition of autophagy and promotion of apoptosis, because loss of LC3 ubiquitination did the opposite as shown in this study (fig. S6D). We further show that LC3 most likely binds the LIR motif of BIRC6 (4670-NPQ TSS <u>F</u>LQ<u>V</u> LV-4681), which does not overlap with the binding regions of Smac, following dimer disruption of this BIRC6 region, providing a basis for the coordinated regulation of apoptosis and autophagy. To remove BIRC6, the ubiquitin ligase Nrdp1 promotes ubiquitination and proteasomal degradation of BIRC6 in response to apoptotic stimuli as we showed previously (*10*). Accordingly, Nrdp1 was also found in the BIRC6 complex (fig. S1E). Given that mutation or overexpression of BIRC6 is frequently associated with various cancers (*32–34*), our findings might contribute to the discovery of therapeutic targets for the enhancement or inhibition of autophagy and apoptosis.

## MATERIALS AND METHODS

### Purification of BIRC6

The plasmid pCI-Myc-BIRC6 (mouse) was a gift from Dr. Stefan Jentsch as described(*6*). A 3×FLAG tag was then inserted into the plasmid ahead of the N-terminus of BIRC6 gene. The plasmids were then transiently transfected into HEK293F expression system for overexpression. After a 48-hrs incubation, the cells were collected, washed one with ice-cold 1×PBS, and then resuspended in buffer A (20 mM HEPES-KOH, pH 7.5, 10 mM KCl, 1.5 mM MgCl_2_, 1mM EDTA, 1 mM EGTA, 1 mM DTT, 10% glycerol, 0.1 mM PMSF, 10 mM Disodium β-glycerophosphate pentahydrate, 2.5 mM sodium pyrophosphate, 1mM NaF, 1 mM Na_3_VO_4_, 1 μg/ml Benzonase Nuclease and a protease inhibitor mixture). The resuspended cells were then lysed by sonication, and the lysate supernatant after centrifugation was incubated with M2-FLAG affinity beads for 1.5 hours at 4°C. After several round of washing with buffer B (20 mM Tris-HCl, pH 8.0, 500 mM NaCl, 10% glycerol, 1 mM ZnCl2, 1 mM NaF, 1 mM Na3VO4 and the protease inhibitor mixture), the FLAG-affinity proteins were eluted by 0.5 mg/ml 3×FLAG peptide (MDYKDHDGDYKDHDIDYKDDDDK) dissolved in buffer C [Phosphate-buffered saline (PBS) supplemented with 100 mM KCl, pH7.4]. The elute was further purified by gel filtration (Superose 6 Increase Columns, Cytiva) using buffer C. To optimize the cry-EM grids preparation for data collection, the peak fractions of BIRC6 were gathered and incubated with 0.01% glutaraldehyde for 10 min, which was then quenched by 40 mM Tris-HCl, pH7.4 buffer. The cross-linked sample was then purified by another round of gel filtration. The peak fractions were pooled and concentrated for cryo-EM grid preparation.

### Purification of His-tagged proteins

1 mM IPTG was used to induce the expression of His-Caspase9-4M (E306A–D315A–D330A-C287A)-pET30a and His-LC3B-pET-30a in BL21 for overnight at 16 °C. After a 16-hrs incubation, the cells were collected, washed once with ice-cold PBS. Lysis /binding /wash buffer contained 300 mM NaCl, 25 mM HEPES, 20 mM Imidazole, and 1μg/ml Benzonase, pH 7.5. The resuspended cells were then lysed by sonication, and the lysate supernatant after centrifugation was incubated with Ni-NTA beads for 20 minutes at 4°C. The His-tagged proteins were eluted by the elution buffer containing 300 mM NaCl, 25 mM HEPES, and 250 mM Imidazole, pH 7.5. The elute was further purified by gel filtration (Superdex TM 75 10/300 GL Columns, Cytiva) using buffer C (PBS supplemented with 100 mM KCl, pH7.4).

### In vitro BIRC6 binding assays

FLAG-BIRC6 was incubated with about 0.5 mg His-Caspase9-4M-pET30a with the molar ratio at 1:300 for 30 minutes at RT. About 0.5mg His-LC3B-pET-30a was incubated with FLAG-BIRC6 in the absence or presence of 100 μM BIRC6 LIR1 (NPQTSSFLQVLV) for 30 minutes at RT. The molar ratio between FLAG-BIRC6 and His-LC3B-pET-30a was 1:300. The mixture was then purified by another round of gel filtration (Superose 6 Increase Columns, Cytiva).

### Electron microscopy

The quality of the purified BIRC6 samples were examined by negative staining EM. In brief, 2-4 μl aliquots of samples were applied on the glow discharged copper grids, and blotted by filter papers after 30-60s’ waiting time. The samples were then washed and stained by 2% (w/v) uranyl acetate solution. The grids were screened using a JEOL JEM-1400Flash microscope operated at an accelerating voltage of 80 kV.

To prepare cryo-EM grids, 3-4 μl aliquots of samples were applied on the glow discharged holey carbon gold grids (C-flat, 2/2, 300 mesh), and flashed frozen in liquid ethane after blotting, using an FEI Vitrobot IV with 100% humidity and 4 °C. The grids were then screened using an FEI Talos Arctica microscope operated at 200 kV. The qualified grids were recovered and transferred into a 300-kV FEI Titan Krios microscope for data collection. The dataset was auto-collected using the SerialEM software (*35*), at a nominal magnification of 64,000 X, corresponding a calibrated pixel size of 1.37 Å at object scale, with a defocus rang of −2 to −3 μm. The images were recorded using a Gatan GIF K3 camera at a super-resolution counting mode with a dose rate of 17.8 e-/s/Å2 and a total exposure time of 3.84s, and at a movie recording mode with 32 frames for each micrograph.

### Image processing

7331 movie stacks were collected for the BIRC6 samples and pre-preprocessed in RELION4.0 (*36*). Local drift correction, electron dose weighting and 2-fold binning were applied to the super-resolution movie stacks using MotionCor2, generating the summed micrographs in both dose-weighted and not dose-weighted versions (*37*). The unweighted micrographs were used in the steps of CTF estimation with CTFFIND4 (*38*), particle picking and initial classification, while the dose-weighted micrographs were then used for fine 3D classification and final refinement.

Around 1000 micrographs were first selected and subjected to several rounds of manual and auto-particle picking, 2D and 3D classifications, using both RELION and CryoSPARC (*39*), to produce the accurate 2D averages for auto-particle picking. With these 2D averages, template matching auto-picking was applied to the complete dataset, with 1,925K particles picked. After two rounds of 3D classification, 72K particles were kept, with their 3D density maps showing fine structural features (fig. S2C). To recover more particles from the the dataset, Topaz was further used to pick particles employing the 72K good particles as the training dataset, resulting in a set of 1583K particles (fig. S2D) (*40*). Similarly, after two rounds of 3D classification, 95K good particles were kept. To avoid that some good particles were mistakenly sorted into the discarded groups during classification, a second round of 3D classification for both particles sets from template matching and Topaz auto-picking were repeated, but in a supervised method, and with the two sets of density maps from the two particle sets serving as the references in a reciprocal manner (fig. S2, C to D). Finally, after merging the four sets of good particles and removing duplicates, a final set of 154K particles were subjected to CTF refinement, bayesian polishing and C2-symmetry imposed 3D refinement, generating a final global density map at 3.6 Å resolution. The resolution of the BIRC6 core region was improved to 3.5 Å by a round of focused refinement (fig. S2F).

Since the local resolutions of the two ends of ArmRD were relatively lower than the central region, the density maps were further refined focused on an half of the U-shaped structure with C2 symmetry expanded particles (fig. S2G). As a result, a significant improvement in structural details was observed for both the ends of ArmRD. On the basis of these optimized density maps, the local density of the N-terminal region was further optimized by a skip-alignment focused 3D classification, which showed the dynamic nature of the N-terminal region (fig. S2H). One group of particles (45K) whose local density map presented improved structural details, was subjected to further refinement (fig. S2H).

The residual density in the central cavity was also optimized on the basis of the global map to identify potential substrate. With C2 symmetry expanded particles, a skip-alignment focused 3D classification separated two groups of particles with Smac docked in the central cavity. In one group, Smac leans to the left side of the cavity while in another group Smac leans to the right side (fig. S2I). The binding modes of Smac to BIRC6 in the structures of the two groups were similar, thus their sorting into two groups might be due to the global C2 symmetry of the BIRC6 dimer. To merge the two groups of particles, these particles were further processed by a round of supervised 3D classification after duplicate removal and C2 symmetry expansion (fig. S2I). All the Smac-bound particles were then subjected to a global refinement, yielding a density map at 4.7 Å resolution. In the groups of particles with other than Smac, residual densities could also be found in the central cavity (fig. S2I). Accordingly, some other substrates of BIRC6, including caspases 3, 6, 7 and 14, HtrA2 and LC3B, and interacting proteins, such as UBA6 and Nrdp1, were also detected in the MS analysis (fig. S1D). But these densities couldn’t be well assigned due to the resolution limit.

### Model building

The initial model for model building was predicted using Alphafold2 (*41*). The sequence of mouse BIRC6 was divided into two overlapped regions (residues 1-2588 and residues 1011-4882), and subjected to model prediction using locally installed Alphafold2. The predicted models were separated into domains or secondary structural motifs, and manually fitted into the density map using Chimera (*42*). Model adjustment or rebuilding was next performed using COOT (*43*). The model was further refined using phenix.real_space_refinement (*44*) with secondary structural and geometry constraints applied.

### Cell Culture

HEK293F cells were cultured in HEK293 Cell Complete Medium Animal Free, Serum Free (Sino Biological). And the HEK293F cells were kept at 37°C with 5% CO_2_ and incubation for 120 rpm. MDA-MB-453, 293T, HeLa and COS7 cells were cultured in Dulbecco’s modified Eagle’s medium (DMEM) supplemented with 10% fetal bovine serum (FBS), 100 U/ml penicillin and 100 μg/ml streptomycin. Wild-type and the Atg5-deficient mouse embryonic fibroblast (MEF) cells were kindly provided by Profs. Y. Ohsumi and A. L. Goldberg. MEF cells were cultured in the above medium with addition of 1% non-essential amino acids and 200 μM β-mercaptoethanol. All cells were kept at 37°C with 5% CO_2_. Cell starvation was performed by incubating cells in HBSS medium at 37°C with 5% CO_2_ after the PBS wash for twice. Rapamycin (Sigma-Aldrich) was used in the normal DMEM medium at the indicated concentration for 24 h. Bafilomycin A1 (Enzo life sciences) was used at a final concentration of 10 nM in HBSS medium for the time points indicated.

### Plasmids and transfection

Myc-tagged Smac plasmids were constructed as described previously (*13*). Myc or FLAG-tagged procaspase-9 plasmid was prepared by cloning the procaspase-9 gene into a pcDNA-based vector (Invitrogen). Procaspase-9 with three mutations (Asp315Ala, Asp330Ala, and Glu306Ala; Procasp9-3M) or with 4 mutations (E306A–D315A–D330A-C287A; Procasp9-4M) was generated using a direct mutagenesis strategy. 293T cells were transfected with plasmids using Lipofectamine 2000 according to the manufacturer’s protocol (Life Technologies, Inc.)

### Confocal immunofluorescence microscopy

Cells grown on glass coverslips in 6-wells were fixed in 4% polyformaldehyde for 10 min at room temperature, followed by membrane permeation using 100 μg/ml digitonin in PBS for 5 min. Cells were blocked in 3% BSA before primary antibodies were applied. Anti-procaspase-9 mouse IgG (1:100, Santa Cruz) and anti-LC3 rabbit IgG (1:100, MBL) were incubated in a moist container for 1 h at room temperature. Then the secondary anti-mouse antibodies conjugated with FITC and anti-rabbit with Alex-594 were applied to the cells in a moist container for 1 h at room temperature. The DNAs in nuclei were stained with 4’-6-diamidino-2-phenylindole (DAPI) at the final preparation step. The slides were viewed on a Zeiss LSM 700 confocal microscope using a 100×oil objective and laser lines at 488 nm and 555 nm for excitation.

### Apoptosis assays

Apoptotic DNA fragmentation was analyzed using the DeadEnd^TM^ Fluorometric TUNEL System according to the standard paraffin-embedded tissue section protocol (Promega). The DNAs in nuclei were stained with DAPI at the final preparation step. The slides were viewed on a Zeiss LSM 700 confocal microscope using a 100 ×oil objective and laser at 488, 555, or 639 nm for excitation.

### Immunoblotting and antibodies

The cell lysis for caspases was carried by sonication in lysis buffer (20 mM Hepes-KOH, pH 7.5, 10 mM KCl, 1.5 mM MgCl_2_, 1 mM EDTA, 1 mM EGTA, 1 mM DTT) with the protease inhibitor mixture (Roche) after treatment by starvation in the HBSS starvation medium with 1 g/l glucose, but without serum or amino acids, for up to 6 h or by rapamycin. For other proteins, the lysis buffer contained 20 mM Tris.Cl, pH 8.0, 500 mM NaCl, 1% NP-40, 10% glycerol, and 1 mM ZnCl_2_. Equal amounts of protein samples were separated by SDS-PAGE and transferred onto PVDF membranes. Antibodies or antisera against the BIRC6 (1:1000, BD Biosciences, #611193,) XIAP (1:1000, BD Biosciences, #610717), Smac (1:1000, Zymed, #35-6600), HtrA2 (1:1000, oncogene, #Am82), ATG5 (1:1000, Cell signaling technology, #12994), Myc (1:1000, Santa Cruz, #B0116), GFP (1:3000, Absin, #137960), Apaf-1 (1:1000, Cell Signaling Technology, #8969), PA28γ (1:3000, Enzo, #PW8190), Anti-GAPDH (1:3000, Santa Cruz, # 25778), p62 (1:1000, Cell signaling technology, #88588S), FLAG(1:3000, Sigma-Aldrich, #F3165), Caspase-9 (1:1000, Cell Signaling Technology, #9508), Cleaved Caspase-3 (1:1000, abcam, ab32042), Caspase-7 (1:1000, Sigma-Aldrich, #SAB4503316), LC3 (1:5000, Sigma-Aldrich, #L7543), β-actin (1:5000, Sigma-Aldrich, #A5441) antibodies were used as primary antibodies to detect the corresponding proteins. Peroxidase-conjugated anti-mouse IgG (1:5000, ZSGB-BIO, #ZB-5305), anti-rat IgG (1:4000, ZSGB-BIO, #ZB-2307) or anti-rabbit IgG (1:3000, ZSGB-BIO, #ZB-5301) was used as the secondary antibody. The protein bands were visualized by an ECL detection system (EMD Millipore).

### Immunoprecipitation

Cells were lysed with the lysis buffer as in the immunoblotting procedure, and the lysates were immunoprecipitated with specific antibody and protein A/G-Sepharose (Sigma-Aldrich) at 4 °C for 2 h. The precipitants were washed 3 times with the lysis buffer, and then eluted with the sample buffer containing 1× SDS at 97 °C for 6 minutes.

### Inhibition of caspase activity assay

To activate caspases in cell lysates, 40 or 80 μg of 293T cell extracts in each reaction (100 μ1) were activated by cytochrome c and dATP essentially as described (*13*). The purified FLAG-tagged BIRC6 was added in the reaction. The caspase activity was analyzed in the buffer containing 0.1 M Tris, pH 6.8, 10 mM dithiothreitol, 0.1% CHAPS, 10% polyethylene glycol, and 20 μM of the caspase 9 substrate Ac-LEHD-amino-4-methylcoumarin (AMC) or caspase 3 substrate Z-DEVD-7-AMC. The release of AMC from the substrates was monitored continuously at 345/445 nm (excitation/emission) at 30 °C for 30 min.

### Quantification and Statistical Analysis

Unless stated otherwise, significance levels for comparisons between 2 groups were determined by one-way ANOVA. Data are reported as mean ±SEM. *P < 0.05; **P < 0.01, ***P < 0.001, normal distribution. All images were chosen at random while blinded, and were quantitated using Image J.

## DATA AND MATERIALS AVAILABILITY

The data that support the findings of this study are available from the authors upon request. The cryo-EM density maps and atomic models have been deposited in the Electron Microscopy Data Bank (EMDB) and Protein Data Bank (PDB): The global density map and atomic model of BIRC6 (EMD-XXXX, PDB: YYYY), the local density map of the N-terminal region (EMD-XXXX), the density map of the Smac-BIRC6 complex (EMD-XXXX).

## ACKNOWLEDGMENTS

We are grateful for Profs. Y. Ohsumi and A. L. Goldberg for kindly providing Atg5^+/+^ and Atg5^-/-^ MEF cells. We thank the cryo-EM platform of Peking University for the cryo-EM grid screening and data collection, the High-performance Computing Platform of Peking University for the computation, and the National Centre for Protein Sciences at Peking University for negative-staining EM and mass-spectrometry analyses. This study was supported by the National Key R & D Program of China (2019YFA0802100 to X.B.Q., 2019YFA0508904 to N.G.), the National Science Foundation of China (31922036 and 32271257 to N.N.L.), and Beijing Municipal Natural Science Foundation (5202014 to T.X.J.).

## AUTHOR CONTRIBUTIONS

S.S.L, T.X.J., F.B., J.L.Z. and G.F.W. devised and performed majority of experiments, and analyzed data with assistance of G.H.Y, J.Y.K, Y.F.Q., L.B.F. and P.W.; X.B.Q., N.G. and N.N.L. conceived the project, supervised the experiments, analyzed data, and wrote the manuscript with input from all authors.

## Supplementary Materials

**Fig. S1.**
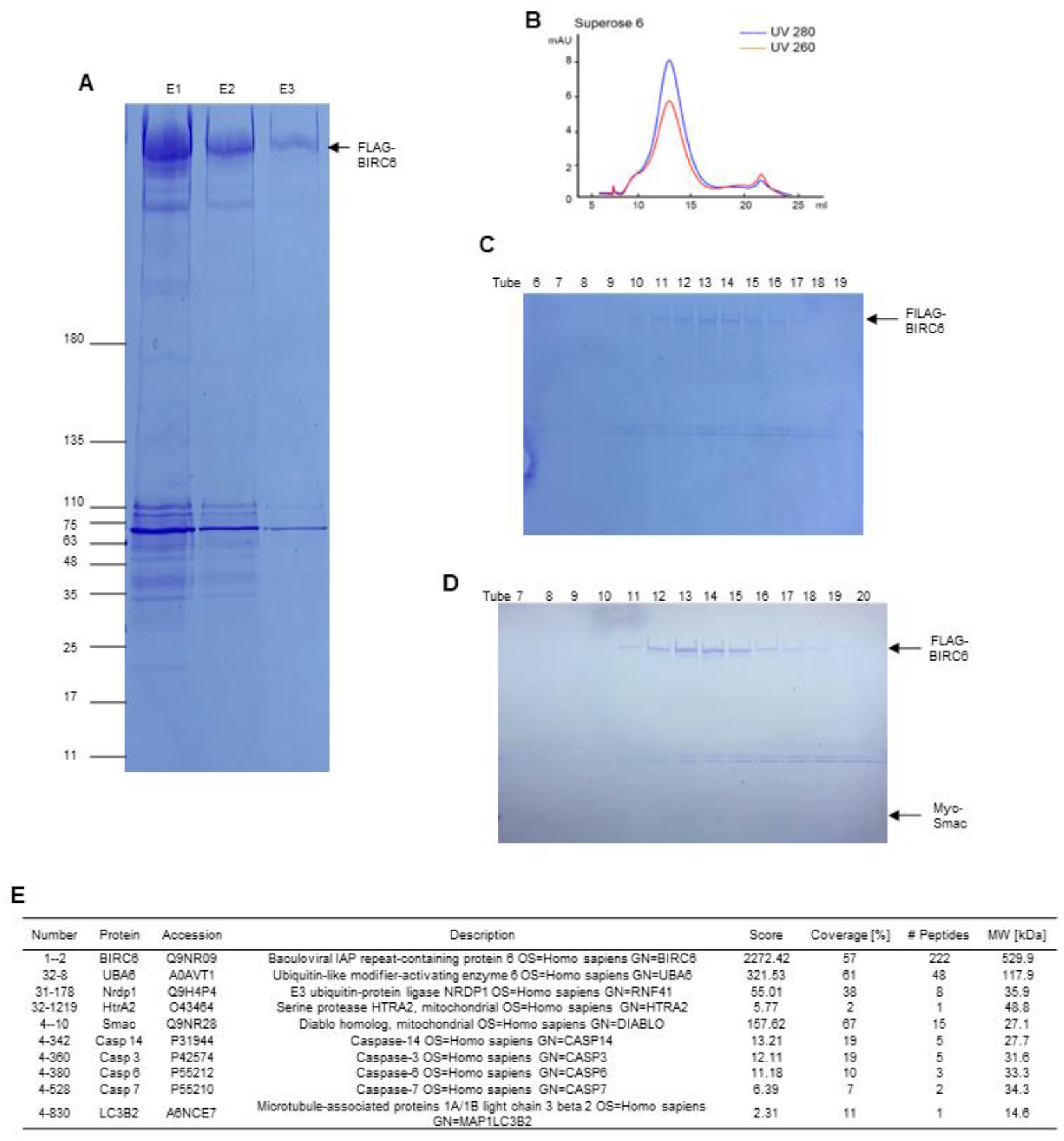
Purification of the FLAG-BIRC6 from HEK293F cells. (**A**) 0.5 mg/ml FLAG peptide in PBS supplemented with 100 mM KCl was used to elute FLAG-BIRC6 at 4°C for 30 min, and repeated for 3 times. Protein samples were analyzed by SDS-PAGE. (**B**) The elutes were passed through the gel filtration-Superose 6 Increase Columns. The peak elution tubes 11-15 were combined for crosslinking 0.01% glutaraldehyde, and passed through the Superose 6 Increase Columns once more. (**C**) The elutes from tubes 6-19 in Fig. S1B were analyzed by Coomassie following SDS-PAGE. (**D**) 0.5 mg/ml FLAG peptide in PBS supplemented with 100 mM KCl was employed to elute FLAG-BIRC6-Myc-Smac complex at 4°C for 30 min, and repeated for 3 times. The elutes were analyzed by Coomassie following SDS-PAGE. (**E**) The purified FLAG-BIRC6 complex was analyzed by mass spectrometry following separation of the complex by SDS-PAGE. The full lane was sliced into 6 pieces from top to bottom with assigned sample numbers: 1, 21, 22, 31, 32, and 4. Potential BIRC6-interacting proteins are listed, and others are provided in online materials.

**Fig. S2.**
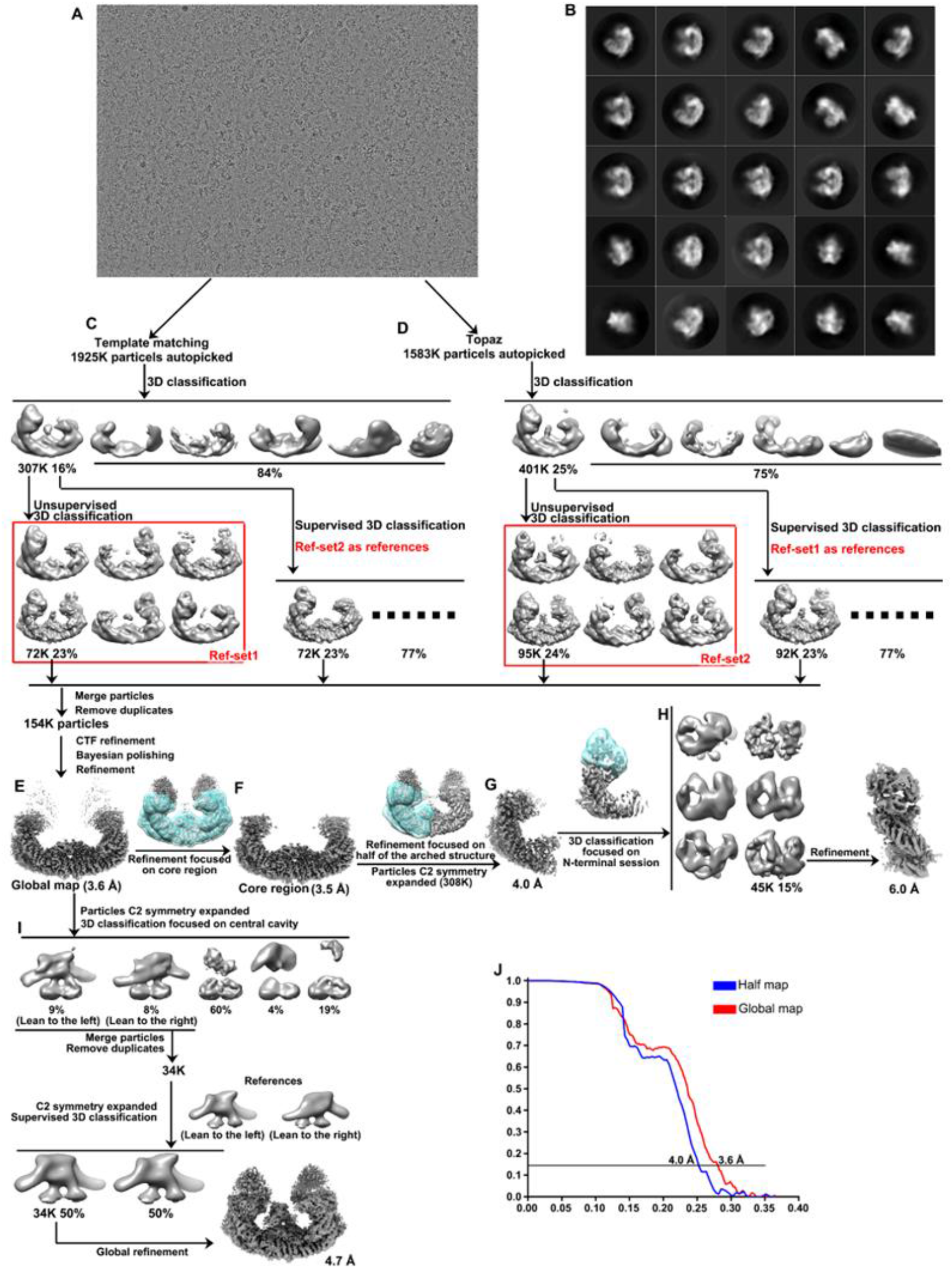
Cryo-EM image processing workflow of BIRC6 sample. (**A**) Representative cryo-EM image of BIRC6 sample. (**B**) 2D class averages of BIRC6. (**C** and **D**) Preliminary 3D classifications of particles autopicked using template matching method Fig. S2C or autopicked using Topaz Fig. S2D. (**E**) The good particles from Fig. S2, C to D were merged and subjected to 3D refinement, generating a final global density map at a resolution of 3.6 Å. (**F**) Refinement focused on the core region. (**G**) Local refinement focused on half of the U-shape structure. (**H**) 3D classification focused on N-terminal session. (**I**) 3D classification focused on the residual density in the central cavity. (**J**) Fourier shell curves from the refinements of global map Fig. S2E or half of the U-shape map Fig. S2G.

**Fig. S3.**
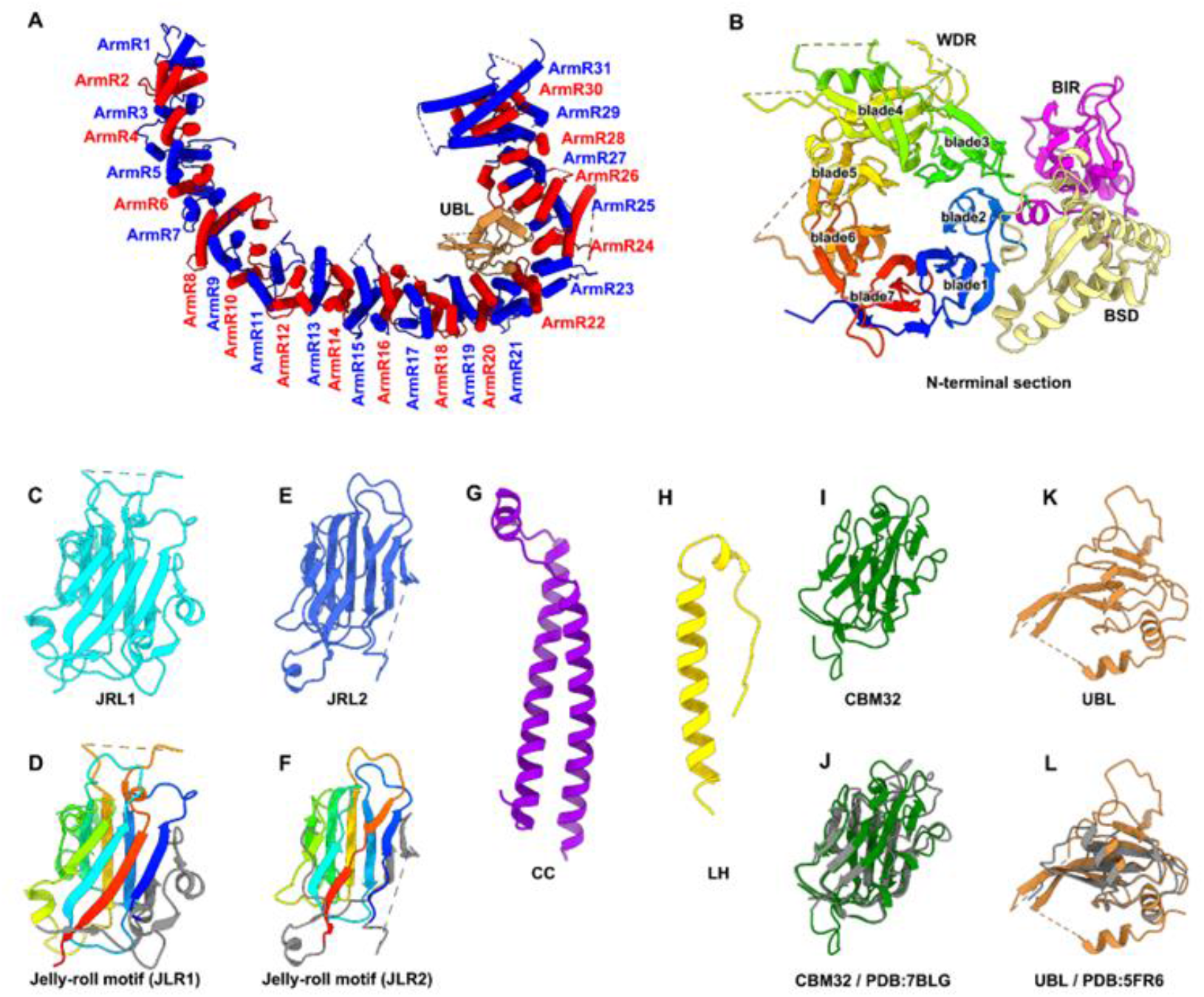
Structures of BIRC6 domains. (**A**) Structure of ArmRD with a sharp bending around ArmR21-24. The 31 ArmRs are colored blue and red alternatively. UBL domain located at the bending site is also shown and labeled. (**B**) Structure of the N-terminal region. WDR is colored rainbow and the seven blades are labeled. BIR and BSD are colored-colored as that in Fig. 1, (C) Structure of JLR1. (**D**) Same as Fig. S3C, but with the jelly-roll motif colored rainbow and the left sequences colored gray. (**E** and **F**) Similar to Fig. S3, C to D, but showing the structure of JLR2. (**G**) Structure of CC. (**H**) Structure of LH. (**I**) Structure of CBM32. (**J**) Same as Fig. S3I, but with another CBM32 structure (PDB: 7BLG) superimposed. (**K**) Structure of UBL. (**L**) Same as Fig. S3K, but with a ubiquitin structure (PDB: 5FR6) superimposed.

**Fig. S4.**
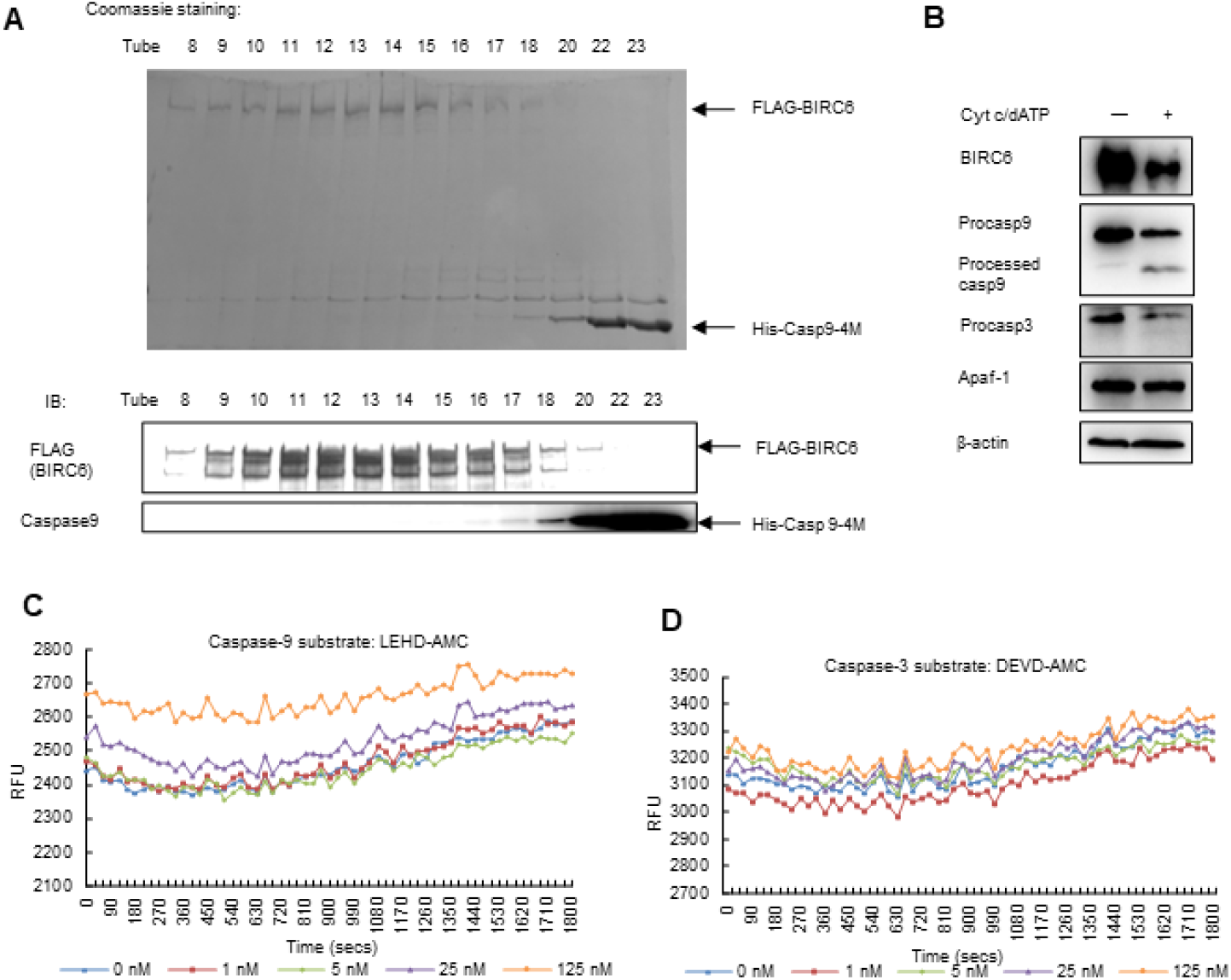
Purified BIRC6 strongly inhibits the activity of caspase 9, but only weakly for caspase 3. (**A**) Caspases in HEK293T cell extracts were activated by cytochrome c and dATP, and protein levels were analyzed by immunoblotting. (**B** and **C**) Active caspases in HEK293T cell extracts were incubated with the purified FLAG-tagged BIRC6 at indicated concentrations. The caspase activities were analyzed using the caspase 9 substrate Ac-LEHD-AMC Fig. S4B or caspase 3 substrate Z-DEVD-7-AMC Fig. S4C. The release of AMC from the substrates was monitored continuously at 380/460 nm (excitation/emission) at 30 °C for 60 min. (**D**) About 0.5 mg His-procasp9-4M was added to 1.2 ml of FLAG-BIRC6 at 0.03 μM in elution buffer, and incubated 30 min at 4 °C. The reaction system was passed through a Superose 6 Column equilibrated with PBS plus 150 mM KCl, and tubes 8-18, 20, 22, 23 were collected for Coomassie staining or immunoblotting following SDS-PAGE.

**Fig. S5.**
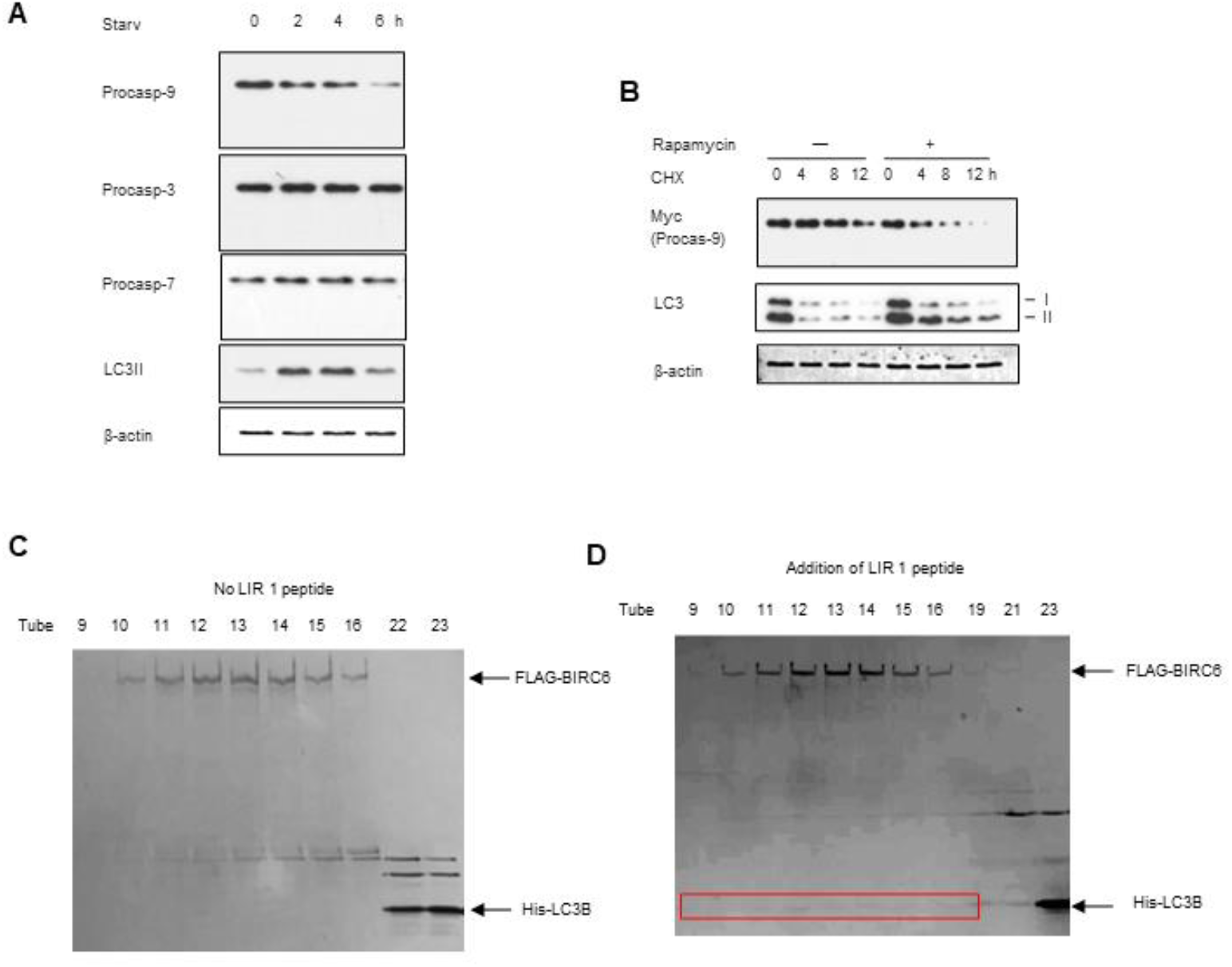
Degradation of Procaspase 9 by autophagy mediated by LC3, whose binding to BIRC6 is enhanced by LIR1 peptide. (**A**) MDA-MB-453 cells were starved in HBSS for 0, 2, 4, or 6 h. The protein levels were detected by immunoblotting. (**B**) HEK293T cells were transfected with Myc-tagged procasp9-3M for 24 h, and then treated with 0 or 0.5 μM rapamycin. After rapamycin treatment for 24 h, 50 μg/ml CHX was added and incubated for the indicated periods of time. The protein levels were detected by immunoblotting. Data are representative of one experiment with three independent biological replicates. (**C** and **D**) About 0.5 mg of His-LC3B were incubated with FLAG-BIRC6 at 0.1 μM in 1.2ml of elution buffer in the absence Fig. S5C or the presence Fig. S5D of LIR1 peptides at 4 °C for 30 min. The reaction system was passed through a Superose 6 Column equilibrated with PBS plus 150 mM KCl, and tubes 8-18, 20, 22, 23 were collected for Coomassie staining following SDS-PAGE. His-LC3 was only co-eluted with FLAG-BIRC6 in the presence of LIR1 peptides (faint bands in the red frame).

**Fig. S6.**
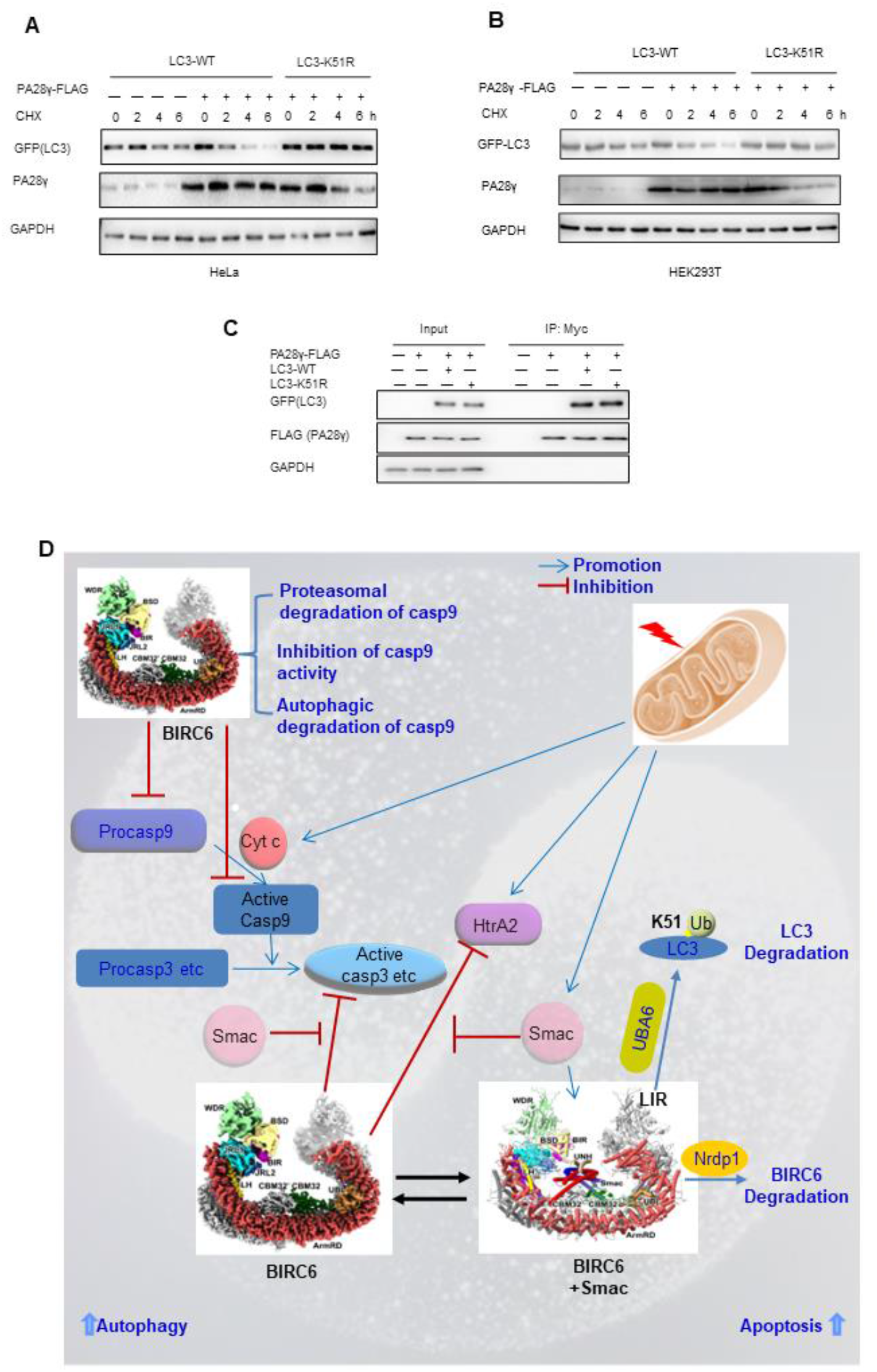
Deficiency in LC3 ubiquitination promotes autophagy, but suppresses apoptosis. (**A** and **B**) HeLa cells (A) or HEK293T (B) cells were co-transfected with GFP-LC3-WT or LC3-K51R and PA28γ-FLAG, and then treated with 100 μg/mL CHX for the indicated periods of time. Protein levels were analyzed by immunoblotting, and data are the results of three independent repeated experiments. (**C**) HEK293T cells were co-transfected with GFP-LC3-WT or LC3-K51R and PA28γ-FLAG. Protein levels were analyzed by immunoblotting following coimmunoprecipitation using the anti-FLAG antibody. (**D**) Model mechanisms by which BIRC6 balances apoptosis and autophagy. A background of Taiji diagram hints the nature of the balance between apoptosis and autophagy.

**Table S1.** Cryo-EM data collection, refinement and validation statistics of Triton X-100 treated sample.

|  | Global map of BIRC6<br>(EMDB-xxxx)<br>(PDB xxxx) |
| --- | --- |
| **Data collection and processing** |  |
| Nominal magnification | 64,000x |
| Voltage (kV) | 300 |
| Electron exposure (e-/Å<sup>2</sup>) | 68.4 |
| Defocus range (µm) | -2 to -3 |
| Pixel size (Å) | 1.37 |
| Micrographs | 7331 |
| Symmetry imposed | C2 |
| Initial particle images | 1925K/1583K |
| Final particle images (total) | 154K |
| Map resolution (total) (Å) | 3.6 |
| FSC threshold | 0.143 |
| **Refinement** |  |
| Model composition |  |
| Non-hydrogen atoms | 47600 |
| Protein residues | 6176 |
| R.m.s. deviations |  |
| Bond lengths (Å) | 0.01 |
| Bond angles (°) | 1.75 |
| Validation |  |
| MolProbity score | 1.68 |
| Clashscore | 5.95 |
| Poor rotamers (%) | 0.00 |
| Ramachandran plot |  |
| Favored (%) | 94.96 |
| Allowed (%) | 4.95 |
| Disallowed (%) | 0.10 |

